# Transcription-based dissection of floral identity and trichome biosynthesis pathways in *Cannabis sativa* L

**DOI:** 10.1101/2025.09.22.677808

**Authors:** Kevelin Barbosa-Xavier, Thiago M. Venancio

## Abstract

*Cannabis sativa* L. has a long history of medicinal and industrial use, with female flowers being the primary source of bioactive compounds such as cannabinoids and terpenoids, which are synthesized in glandular trichomes. Despite its importance, the genetic mechanisms governing flower development, sex determination, and the biosynthesis of these valuable secondary metabolites remain only partially understood. In this study, we conducted an in-depth transcriptomic and comparative genomic analysis to elucidate the molecular networks underlying these key traits. By integrating 117 RNA-Seq datasets and performing phylogenetic analyses across nine distinct *C. sativa* genomes, we identified 31 orthogroups of MADS-box transcription factors. Expression profiling of these genes revealed distinct sets of candidates associated with male and female flower identity, consistent with the established ABCDE model of floral development. Specifically, genes from the AP3, PI/GLO, and MIKCS clades showed preferential expression in male flowers, while genes from the AGL6, FLC-like, and Bsister clades were highly expressed in female flowers. Furthermore, our investigation into genes related to pollen development highlighted the significant role of sugar metabolism and transport in male fertility. Finally, our analysis of cannabinoid and terpenoid biosynthetic genes confirmed their pronounced expression in trichomes and highlighted a key distinction: while the upstream polyketide, MEV, and MEP pathways and the terpenoid pathway showed conserved expression across chemotypes, the cannabinoid pathway exhibited chemotype-specific expression profiles. Collectively, our findings provide a comprehensive molecular framework for understanding floral development and secondary metabolism in *C. sativa*, offering valuable targets for future functional studies and advanced breeding programs aimed at optimizing desirable agronomic and medicinal traits.

## Introduction

The use of *Cannabis sativa* L. female flowers (FF) dates back over 2,700 years, serving as medicinal, ritualistic, and recreational purposes across diverse cultures (Ren et al., 2019). In modern contexts, cannabis is primarily consumed via two methods: combustion (smoking) and essential oil extraction. FF are the most valuable part of the plant due to their rich biochemical profile (Pieracci et al., 2021; Ren et al., 2019). These flowers are densely covered in glandular trichomes—microscopic resin-producing structures that synthesize a wide array of secondary metabolites, including cannabinoids, terpenoids, and flavonoids. These compounds are responsible for the plant therapeutic, psychoactive, and sensory properties (ElSohly & Slade, 2005; Ren et al., 2019). The primary metabolites produced in cannabis FF are cannabinoids, hydrocarbons, sugars, steroids, mono- and sesquiterpenes, flavonoids, amino acids, and nitrogenous compounds (ElSohly & Slade, 2005). Among these, cannabinoids are the most pharmacologically and commercially valuable, particularly delta-9-tetrahydrocannabinol (Δ9-THC) and cannabidiol (CBD), which are associated with both medicinal and adult-use applications (Gülck & Møller, 2020; Leinen et al., 2023). Terpenoids also contribute significantly, especially in the context of adult use, as they shape the aroma and flavor profile of cannabis products (Hanuš & Hod, 2020; Rice & Koziel, 2015). Notably, the effects observed from smoking cannabis or using full-spectrum oils result from the combined action of multiple metabolites produced in trichomes, a phenomenon known as “entourage effect” (Ferber et al., 2020). *C. sativa* is a dioecious species, producing separate male and female plants. Although studies have shown that flower sex can be altered by chemical or environmental cues (Adal et al., 2021), the genetic mechanisms underlying flower development and sex differentiation remain largely unresolved. In this study, we leveraged a comprehensive collection of RNA-Seq datasets from the Cannabis Expression Atlas (CannAtlas) (Barbosa-Xavier et al., 2024) to investigate three key gene groups involved in cannabis flower biology: (i) MADS-box transcription factors (TFs); (ii) genes potentially associated with pollen development, and; (iii) genes involved in cannabinoid and terpenoid biosynthetic pathways. These transcriptomic analyses were complemented by comparative genomics across multiple *Cannabis* genomes. Our findings reveal candidate genes associated with flower morphology and sex determination, as well as key enzymes driving cannabinoid and terpenoid production in trichomes.

## Material and methods

### Expression data

Expression data (in transcripts per million, TPM) from 117 RNA-seq samples— comprising trichomes (n = 59), FF (n = 34), male flowers (MF, n = 18), and induced male flowers (IMF, n = 6)—along with their metadata, were obtained from the CannAtlas (Barbosa-Xavier et al., 2024).

### MADS-box phylogenetic and orthology analysis

We used the PlantTFDB v5.0 prediction tool (Tian et al., 2020) to identify MADS-box TFs across nine *C. sativa* varieties with publicly available genome and protein sequence data (Table 1). To determine orthologous relationships across these nine *C. sativa* genomes, we employed OrthoFinder v3.0.1b1 (Emms & Kelly, 2019). The MADS-box protein sequences reported by Ristevski (2023) were used as references to identify orthogroups (OGs) containing MADS-box genes. Only OGs with at least one reference MADS-box protein were retained for downstream analysis. Protein sequences within each OG were aligned using MAFFT v7.525 (Katoh & Standley, 2013), and maximum-likelihood phylogenies were reconstructed using IQ-TREE v2.3.6 (Minh et al., 2020) under default parameters. Phylogenetic trees were visualized with FigTree v1.4.4 (Rambaut, 2012), and OG alignments were further inspected using the NCBI Multiple Sequence Alignment Viewer v1.26.0.

**Table 1.**
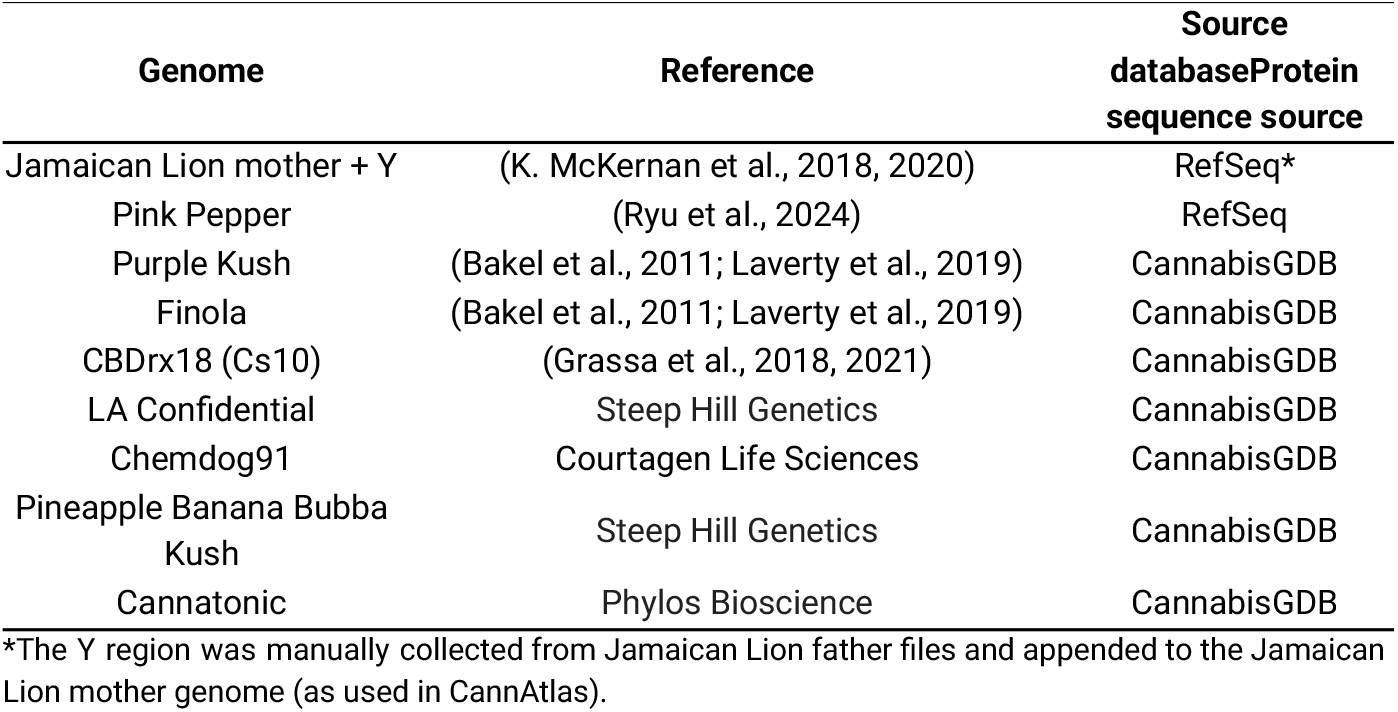
Cannabis varieties used in MADS-box analysis.

### Male flower development genes

We conducted BLAST (Altschul et al., 1997) searches against CannAtlas using three reference gene sets to identify genes potentially related to pollen development and male function in *C. sativa*: (i) 62 sugar metabolism‐related reference genes from Liu et al. (2021), (ii) 61 candidate genes associated with masculinization in cannabis proposed by Adal et al. (2021), and (iii) 185 reference genes implicated in male sterility in Chinese cabbage (*Brassica rapa*) (Huang et al. (2020). BLAST searches were performed with the following parameters: *similarity_matrix* = BLOSUM62, *e-value* ≤ 0.001, *qcov_hsp_perc* ≥80 and, *max_target_seqs* = 1.

### Cannabinoid and terpenoid pathway genes

Genes involved in cannabinoid and terpenoid biosynthetic pathways, as reported by Kovalchuk et al. (2020), were retrieved from Genbank and used to extract expression data from CannAtlas. All genes were successfully identified in CannAtlas. BLAST searches were conducted with the following parameters: *per_identity* = 80, *e-value* ≤ 1e-10, and *qcov_hsp_perc* ≥ 90.

### Enrichment analysis

Enrichment analyses of Gene Ontology (OG), KEGG pathways, and TF family were performed using a Fisher’s exact test in R, with significance defined by a Benjamini-Hochberg adjusted p-value < 0.05.

## Results and discussion

### Identification of MADS-Box orthogroups across cannabis varieties and analysis of their expression in male and female flowers

The MADS-box TF family plays a central role in the ABCDE genetic model of flower identity, which governs the specification and differentiation of sepals, petals, stamens, carpels, and ovules (F. Chen et al., 2017). This model describes how the combinatorial expression of specific MADS-box TFs defines floral organ identity and coordinates flower morphogenesis, ensuring proper reproductive development (Coen & Meyerowitz, 1991). Using *Arabidopsis thaliana* MADS-box TFs as reference, Ristevski (2023) identified 68 cannabis MADS-box TFs in the *C. sativa* ‘Cs10’ genome (CBDrx-18), which were distributed across 18 phylogenetic clades. These include members of both type I (M*α*, M*β*and M*γ*) and type II (MIKCS, Bsister, PI/GLO, AP3, AGL12, SVP, AGL15, AGL17, AG, TM3, FLC-like, FLC, AGL6, SQUA, and SEP) MADS-box genes. To expand this analysis, we used the PlantTFDB prediction tool to identify MADS-box TFs across nine *C. sativa* varieties with publicly available genomes (Table 2). Based on the total number of predicted TFs per genome, we found that MADS-box genes account for an average of 4.57% of all cannabis TFs (Table 2). In the CBDrx-18 (Cs10) genome, our approach identified 76 MADS-box genes, including all the 68 genes reported by Ristevski (2023). This result supports the accuracy and sensitivity of the strategy used in both studies.

**Table 2.** MADS-box TF distribution across nine cannabis varieties.

| Variety | # Total TFs | # MADS TFs | % MADS |
| --- | --- | --- | --- |
| Jamaican Lion mother + Y | 1532 | 70 | 4.57 |
| Pink Pepper | 2315 | 123 | 5.31 |
| Purple Kush | 1816 | 77 | 4.24 |
| Finola | 1694 | 77 | 4.55 |
| CBDrx18 (Cs10) | 1399 | 76 | 5.43 |
| LA Confidential | 1055 | 43 | 4.08 |
| Chemdog91 | 789 | 35 | 4.44 |
| Pineapple Banana Bubba Kush | 920 | 36 | 3.91 |
| Cannatonic | 949 | 44 | 4.64 |
| **Mean** | **1385.44** | **64.56** | **4.57** |

OrthoFinder grouped 95.5% of the input genes (280,293 out of 293,409 predicted proteins) into 17,972 OGs. Among these, 0.5% (1,467 genes) were assigned to 596 variety-specific OGs (Table S1), with the majority concentrated in Purple Kush, Jamaican Lion, and Finola, which together accounted for 420 of the variety-specific OGs (Table S2). To explore MADS-box orthology across *C. sativa* varieties, we used OrthoFinder with the 68 MADS-box TFs identified by Ristevski (2023) in the CBDrx-18 (Cs10) genome as reference. We identified 31 OGs containing at least one MADS-box gene, of which 23 included at least one of the reference MADS-box TFs (Table S3). Among these, only two OGs grouped genes from more than one phylogenetic clade: OG0000096 included representatives of AG, AGL6, FLC, and SEP, while OG0000433 contained genes from the FLC-like and TM3 clades (Table 3 and S3). The remaining reference genes were distributed across distinct OGs as follows: AP3, AGL17, AGL12, AGL15, SVP, SQUA, Bsister, and PI/GLO, each mapped to separate OGs, whereas M*α*, M*β*, M*γ* and MIKCS reference genes were distributed across 5, 3, 3, and 2 OGs, respectively (Table 3 and S3). Some OGs also included genes that were not predicted as MADS-box or as members of any TF family by the PlantTFDB pipeline (Table 3). To understand why these genes appeared in MADS-box clades, we examined their Multiple Sequence Alignments (MSA) and found that, although they showed partial alignment with MADS-box genes, they lost the conserved N-terminal MADS-box domain (Figure 1 and Figures S1–S17). This domain is essential for the canonical function of the MADS-box TFs and its absence explains why the homologs that lost them were not recognized by PlantTFDB. Finally, we used CannAtlas to assess expression patterns of the identified cannabis MADS-box TFs in male and FF (Figure 2).

**Table 3.**
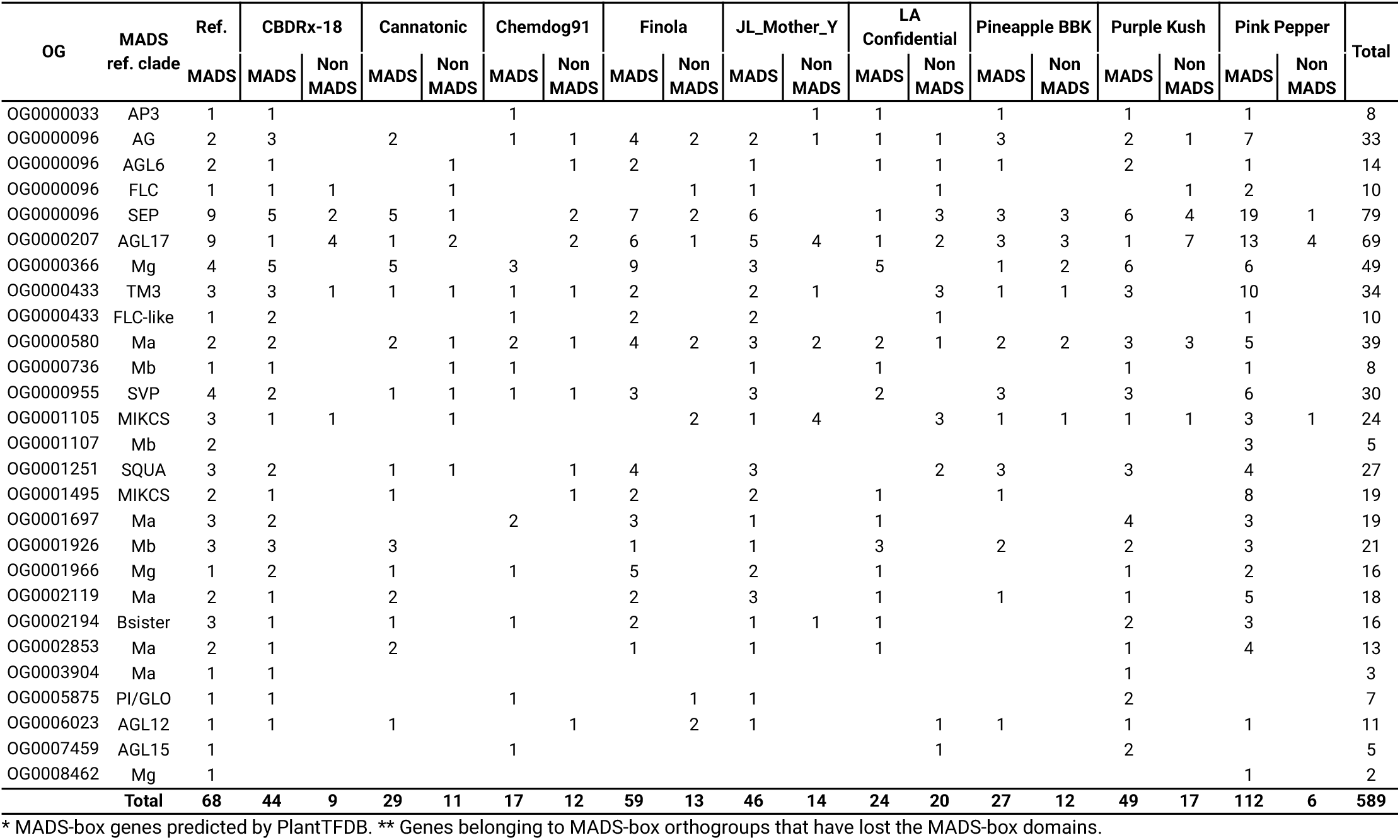
Number of MADS-box* and Non-MADS** genes in each reference clade of.

**Figure 1.**
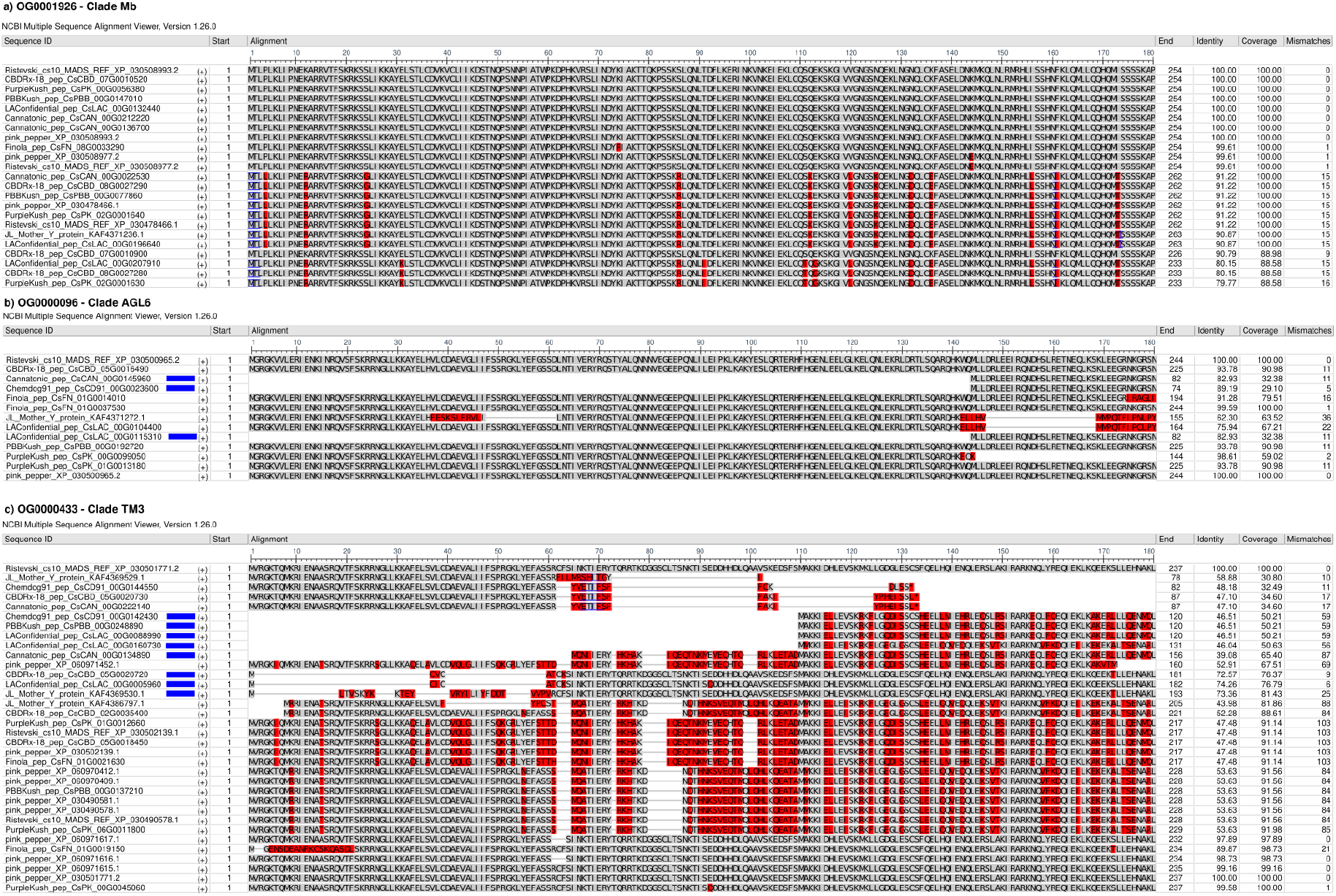
Multiple sequence alignment showing examples of a) a clade composed entirely of genes classified as MADS-box, and two clades containing b) three non-MADS genes and c) eight non-MADS genes. Non-MADS genes are marked with blue rectangles.

**Figure 2.**
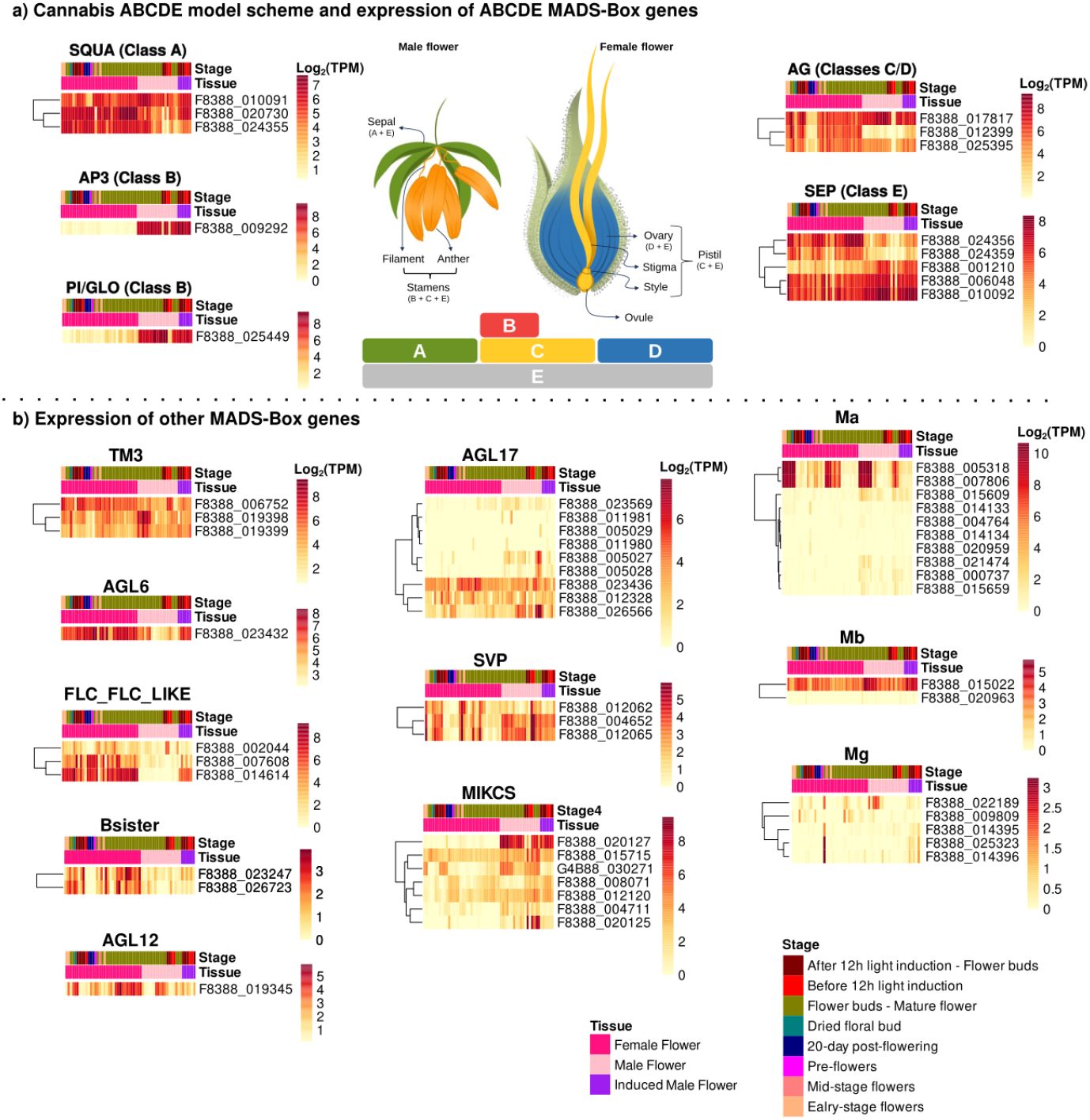
Expression profiles of representative MADS-box genes by clade. Heatmap shows normalized expression for representative genes from each orthology-defined clade. The “Class” column refers to the ABCDE model of floral organ identity, as represented at the central scheme. The “Stage” column indicates the developmental stage for each sample, as reported in the SRA metadata. The central scheme depicts the ABCDE model; colors map classes to organs. Model adapted from (Y.-T. Chen et al., 2019); flower drawings adapted from Releaf (2023), Schilling et al. (2021) and Yadav et al. (2023). Illustration: Rosana Gobbi Vettorazzi.

We identified 60 MADS-box TFs in CannAtlas, representing all clades described by Ristevski (2023), except for AGL15 (Figure 2). Genes from the AGL6, FLC-like, and Bsister clades, as well as SEP genes F8388_024356 and F8388_024359, and the AG gene F8388_012399, showed high expression in FF (Figure 2). In contrast, all genes from the AP3, PI/GLO, SVP, and MIKCS clades, along with the SEP gene F8388_001210, are predominantly expressed in MF (Figure 2). The remaining AG and SEP genes, as well as representatives from other MADS-box clades, exhibit relatively balanced expression between MF and FF (Figure 2). According to the ABCDE model of floral organ identity (F. Chen et al., 2017; Dreni & Kater, 2014; H. Liu et al., 2021), the major classes of MADS-box genes are defined as follows:

- Class A: SQUAMOSA (SQUA) / APETALA1 (AP1);
- Class B: APETALA3 (AP3) and PISTILLATA (PI/GLO);
- Class C: AGAMOUS (AG);
- Class D: SHATTERPROOF (SHP) and SEEDSTICK (STK), (from AG subfamily);
- Class E: SEPALLATA (SEP).

In addition to these core classes, other MADS-box genes such as AGL6, FLC, FLC-like, AGL17, TM3, SVP, MIKCS, AGL12, Bsister, and AGL15 are also implicated in flowering processes (H. Liu et al., 2021; Ristevski, 2023; Shah et al., 2022).

The ABCDE model suggests that the male reproductive organ—the stamen (anther and filament)—is specified by the combined action of class B, C, and E genes (Figure 2) (Gioppato & Dornelas, 2019). Based on the expression patterns observed here (Figure 2), we propose that the class B genes AP3 (F8388_009292) and PI/GLO (F8388_025449), the class C/D AG (F8388_017817), and class E SEP (F8388_001210) are strong candidates for regulators of MF identity in *C. sativa*. Additionally, the MIKCS genes—particularly F8388_020127—also show higher expression in MF, suggesting a possible role in MF development (Figure 2). Although MIKCS genes have been previously linked to pollen development (Gramzow & Theissen, 2010), their roles in broader aspects of floral morphogenesis remain poorly understood. MADS-box genes from classes C, D, E, and Bsister are known to be involved in the development of female reproductive organs such as the ovary, style, and stigma (Figure 2) (de Folter et al., 2006; Gioppato & Dornelas, 2019). Our data support that the C/D gene AG (F8388_012399), the E genes SEP (F8388_024356 and F8388_024359), and Bsister genes (F8388_023247 and F8388_026723) are likely regulators of FF identity in cannabis (Figure 2). In addition, AGL6 (F8388_023432) and FLC-like (F8388_007608 and F8388_014614) genes also exhibit higher expression in FF, further suggesting their involvement in FF development (Figure 2). Although experimental validation is required to confirm the functional roles of these candidate genes, our integrative analysis using publicly available transcriptomic data provides a valuable foundation for prioritizing MADS-box TFs involved in sex-specific floral development in *C. sativa*.

### Identification and expression patterns of candidate genes involved in pollen development

To investigate genes beyond the canonical MADS‐box regulators involved in MF development in *C. sativa*, we conducted BLAST searches against CannAtlas using three reference gene sets. First, from the 62 sugar metabolism‐related genes reported in maize by Liu et al. (2021), we identified 35 homologs in CannAtlas, of which 10 displayed preferential expression in both genetic MF and IMF (Figure 3, Table S4). Second, among the 61 candidate genes proposed by Adal et al. (2021) as associated with masculinization in cannabis, 37 homologs were detected in CannAtlas, with 13 showing preferential expression in MF and IMF (Figure 3, Table S4). Third, from the 185 reference genes implicated in male sterility in Chinese cabbage by Huang et al. (2020), we identified 42 homologs in CannAtlas, of which 23 exhibited preferential expression in MF and IMF (Figure 3, Table S4).

**Figure 3.**
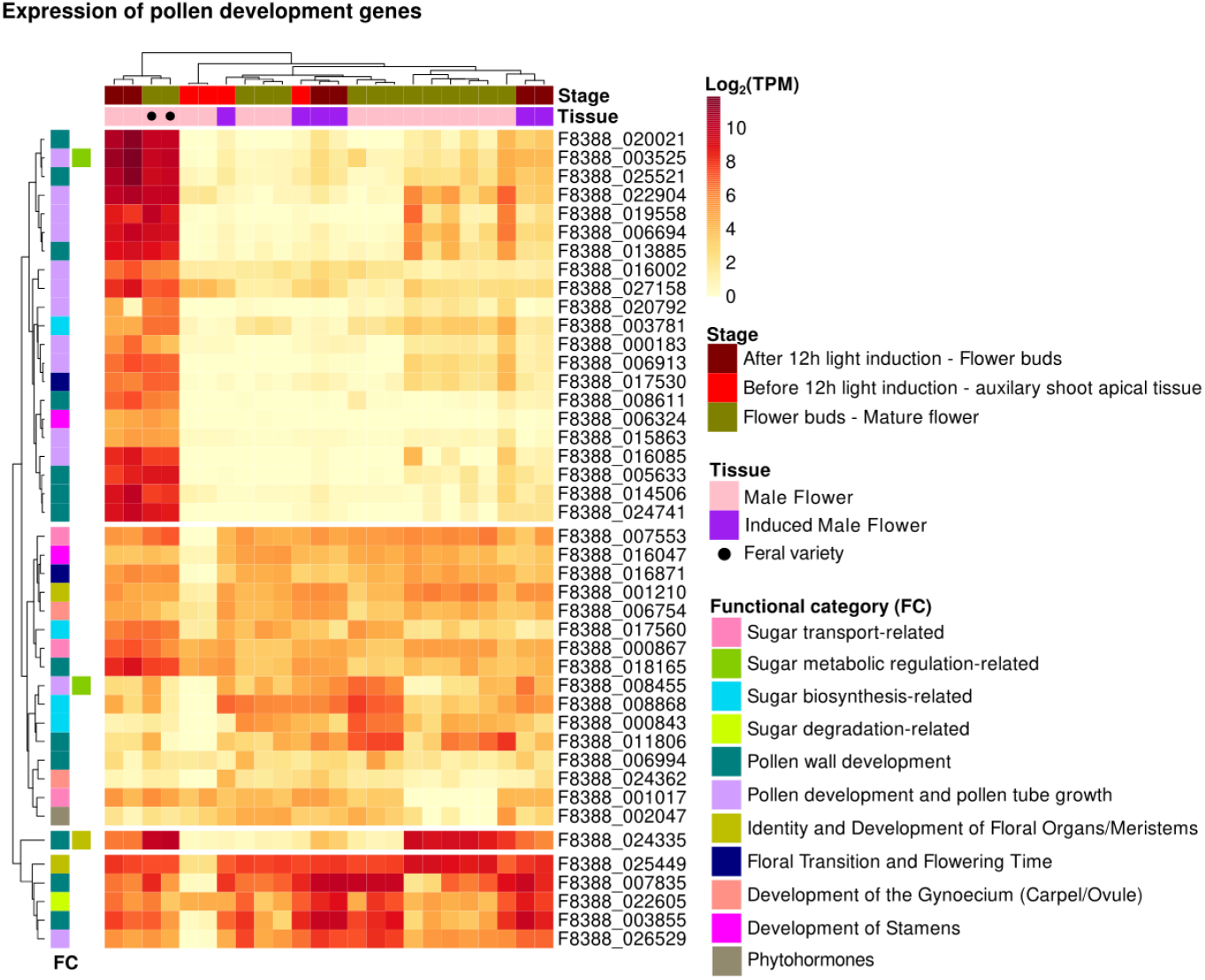
Expression patterns of genes associated with pollen development, classified into functional categories (FCs). “Stage” refers to the developmental stage of the samples, as reported in the SRA database.

Huang et al. (2020) grouped their gene set into four functional categories: pollen development and tube growth, pollen wall development, phytohormone regulation, and TFs. Similarly, Adal et al. (2021) classified their candidates by specific functions in flower development. Building on these frameworks, we assigned the 43 CannAtlas genes identified in our analysis to 11 functional categories (FC) related to flower and pollen development (Figure 3, Table S5).

Notably, nearly all genes under “pollen development and pollen tube growth”, together with approximately half of those in “pollen wall development”, exhibit high expression in MF samples collected after 12 hours of light induction—coinciding with the onset of floral development. A similar expression pattern was also observed in MF samples from two feral varieties (SAMN20863695 and SAMN20863694). In contrast, 22 genes across multiple categories show comparable expression levels in both MF and IMF, with most showing little to no expression in MF samples collected before the 12-hour light induction mark (i.e., at the axillary shoot apical meristem stage). This expression pattern suggests the existence of a light‐triggered developmental switch that initiates a coordinated gene expression program essential for MF development, including genes involved in pollen formation.

Among the 22 genes expressed similarly in MF and IMF were four MADS-box TFs. Two of these—F8388_001210 (class E, SEP) and F8388_025449 (class B, PI/GLO)—are canonical ABCDE model genes that govern floral organ identity and are essential for stamen specification. The other two—F8388_006754 and F8388_016871—are non-ABCDE MADS-box genes. F8388_006754 has been associated with gynoecium development, while F8388_016871 has been implicated in floral transition and regulation of flowering time (Table S5). Interestingly, although F8388_006754 is typically linked to female organ formation, its expression in MF suggests it may play a repressive role in this context. A similar mechanism has been described in *Silene latifolia*, where Hardenack et al. (1994) and Bačovský et al. (2022) showed that altered expression of MADS-box genes, including class B genes PISTILATA and APETALA3, correlates with repression of gynoecium development in MF. Likewise, Di Stilio et al. (2005) proposed that in dioecious species, sex determination may involve modulation of floral homeotic gene activity, particularly through the downregulation of female organ identity programs during male flower development. Taken together, these findings support the hypothesis that F8388_006754 contributes to repressing gynoecium formation in male *C. sativa* flowers, thereby facilitating proper stamen development.

Among the 10 sugar-related genes identified, eight displayed similar expression levels in both MF and IMF, suggesting a conserved role in pollen development across genetic and chemically induced contexts. Three of these genes encode sugar transporters: a bidirectional member of the SWEET family (F8388_007553), a sucrose/H^+^symporter (F8388_000867), and a UDP-galactose/UDP-glucose transporter (F8388_001017). These transporters likely help sustain the high energy demands for pollen development and pollen tube elongation. In *Arabidopsis*, disruption of sucrose transport impairs pollen germination and shortens pollen tubes, underscoring the critical role of sugar supply in reproductive success (De Angeli, 2022; Reinders, 2016). In addition, two genes associated with sugar metabolism regulation—a putative L-ascorbate oxidase (F8388_003525) and a MYB TF (F8388_008455)—may contribute to modulating redox balance and secondary metabolism during pollen development, processes that influence osmotic regulation and energy availability, both essential for pollen tube growth (Li et al., 2020). We also identified two hexosyltransferases (F8388_000843 and F8388_008868) and a 1,3-β-glucan synthase (F8388_017560) involved in sugar biosynthesis. Hexosyltransferases catalyze the transfer of hexoses to elongating polysaccharide chains, contributing to the formation of cell wall components such as hemicelluloses and pectins (Li et al., 2020). Meanwhile, 1,3-β-glucan synthases drive callose production, a structural component of the pollen wall that reinforces integrity during maturation (Nishikawa et al., 2005). Collectively, these genes likely coordinate dynamic remodeling of the pollen cell wall through the regulated synthesis and deposition of carbohydrate polymers, ensuring structural stability and functionality in both MF and IMF. Several genes related to pollen wall development also showed comparable expression levels in both MF and IMF, supporting a shared molecular program for pollen wall formation across both flower types. These include a cytochrome P450 (F8388_003855), a NAD-dependent epimerase/dehydratase (F8388_007835), a chalcone synthase

(F8388_011806), a beta-galactosidase (F8388_024335), a GH10 domain-containing protein (F8388_006994), and a pectinesterase (F8388_018165). These enzymes act in the biosynthesis and remodeling of the structurally complex pollen wall layers. Cytochrome P450 enzymes contribute to anther cuticle development and sporopollenin biosynthesis, both essential for the outer exine layer (Ma et al., 2022; Yang et al., 2014). NAD-dependent epimerases/dehydratases facilitate sugar residue modification for polysaccharide biosynthesis (Usadel et al., 2004). Chalcone synthase catalyzes the first committed step in flavonoid biosynthesis, critical for pollen viability and wall integrity (Buer et al., 2010). Beta-galactosidases participate in cell wall remodeling by cleaving galactose-containing polysaccharides, enabling proper wall expansion and maturation (Ban et al., 2020). Pectinesterases modulate pectin methylesterification, thereby adjusting wall porosity, rigidity, and elasticity (Cankar et al., 2014). The GH10-domain protein is likely involved in hemicellulose remodeling, though its role in pollen development remains to be clarified. Additionally, some genes appear to contribute to both pollen development and pollen tube growth, such as a short-chain dehydrogenase/reductase (SDR, F8388_026529) and the previously mentioned MYB TF (F8388_008455), highlight the coordinated regulation of wall biosynthesis and signaling pathways required for male gametophyte function.

### Upstream cannabinoid and terpenoid pathway genes show parallel expression in hemp and marijuana

The FF of *C. sativa* have historically been the most utilized part of the plant due to their psychotropic and medicinal properties. In plant physiology, secondary metabolite production is commonly associated with defense responses against biotic and abiotic stresses. In cannabis, this defensive role is largely mediated by glandular trichomes—specialized epidermal structures on FF that synthesize and store a wide array of secondary metabolites, including cannabinoids and terpenoids. To gain deeper insight into the functional roles of trichome-specific genes identified in CannAtlas (Barbosa-Xavier et al., 2024), we performed enrichment analyses for Gene Ontology (GO), metabolic pathway (KEGG), and TF families. The enriched GO and KEGG terms strongly point to the biosynthesis, modification, and storage of secondary metabolites (Figure 4).

**Figure 4.**
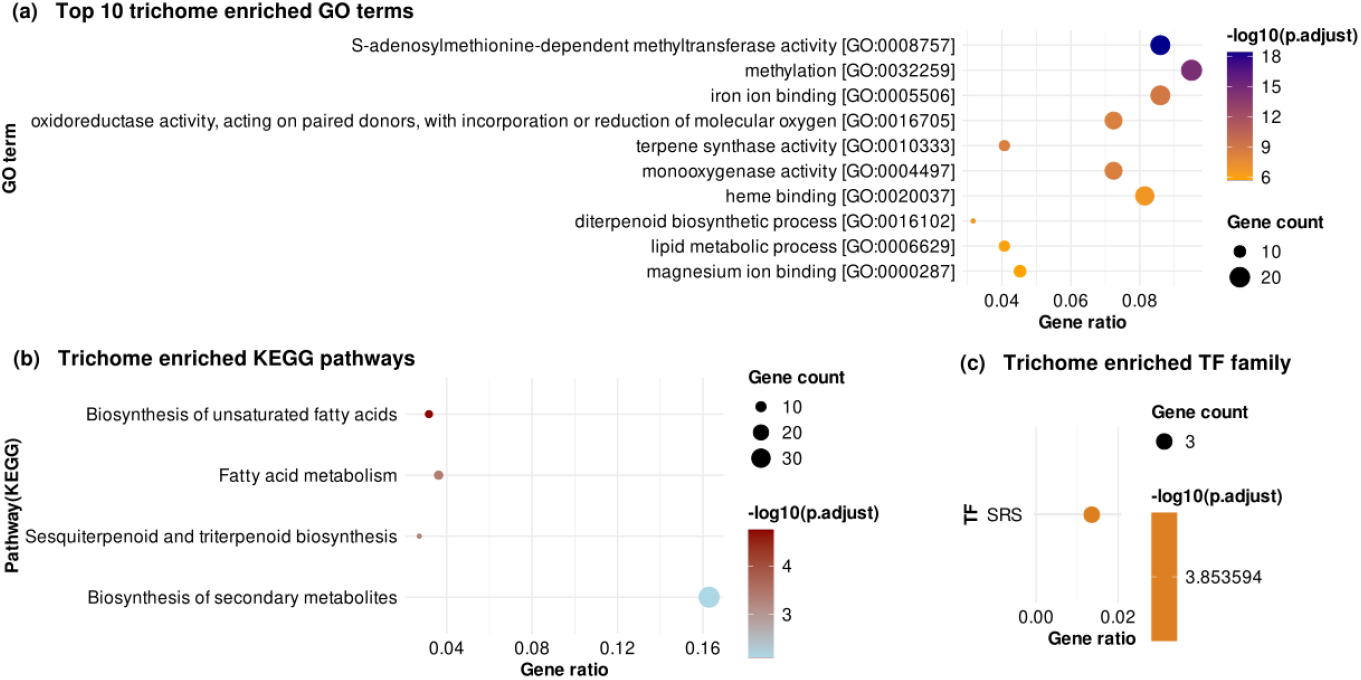
Functional enrichment of trichome-specific genes. (a) Gene ontology (GO) terms, (b) KEGG metabolic pathways and (c) transcription factor (TF) families.

Among the enriched GO terms (Figure 4a), S-adenosylmethionine-dependent methyltransferase activity and methylation are particularly relevant, reflecting the role of methylation in enhancing the bioactivity, chemical stability, and defense potential of secondary metabolites such as cannabinoids and terpenoids (C. Zhang et al., 2021). Additional enrichment in ion binding, heme binding, and monooxygenase activity (Figure 4a) highlights the contribution of cytochrome P450 enzymes, which participate in the structural diversification of secondary metabolites and modulate their chemical properties (Layer et al., 2010). The enrichment of oxidoreductase activity (Figure 4a) underscores the oxidative processes necessary for synthesizing bioactive compounds. Terpene synthase activity (Figure 4a) was also significantly enriched, consistent with the role of volatile terpenes in repelling herbivores and inhibiting microbial pathogens (MacWilliams et al., 2023). GO terms related to the diterpenoid biosynthesis and lipid metabolism (Figure 4a) support the production of cannabinoids and their fatty acid precursors. KEGG enrichment analysis (Figure 4b) corroborated these patterns. Enrichment of unsaturated fatty acid biosynthesis and fatty acid metabolism point to the generation of precursors for signaling molecules like jasmonic acid, a key hormone in plant defense (Luo et al., 2019; Marks et al., 2009). The enrichment of sesquiterpenoid and triterpenoid biosynthesis highlights the trichome role in producing antimicrobial and insect-repellent compounds, while the broad enrichment of secondary metabolite biosynthesis emphasizes the specialized metabolic capacity of trichomes. TF enrichment (Figure 4c) revealed the SRS (SHI-related sequence) family as significantly overrepresented. This TF family regulates both trichome development and secondary metabolism (J. Zhang et al., 2020), suggesting that SRS TFs coordinate structural and chemical components of the cannabis defense system.Taken together, these results highlight cannabis glandular trichomes as hubs of defensive metabolism, integrating gene regulation, specialized metabolite biosynthesis, and adaptive responses to environmental challenges.

Cannabinoids and terpenes are the most extensively investigated compounds in *C. sativa* for their medicinal or recreational importance. A detailed understanding of their metabolic pathways is essential for strategies aimed at manipulating metabolite content in cannabis plants. To identify key genes involved in these pathways, we performed BLAST searches in CannAtlas using proteins previously characterized in cannabinoid and terpenoid biosynthesis (Kovalchuk et al., 2020), including those from upstream pathways such as polyketide, methylerythritol (MEP), and mevalonate (MEV) (Kovalchuk et al., 2020). This search identified 42 genes (Table S6). Given that these secondary metabolites are synthesized and stored in glandular trichomes of FF, we assessed their expression in flower and trichome samples (Figure 5). The polyketide pathway, which produces olivetolic acid, involves three core enzymes: Acyl-Activating Enzymes (AAEs), Olivetol Synthase (OLS), and Olivetolic Acid Cyclase (OAC), with OLS and OAC being directly implicated in cannabinoid biosynthesis (Thomas et al., 2020). We identified 13 polyketide-related genes (Table S6), including six AAEs classified as “expressed-in-all” and two as “mixed” in CannAtlas. Two OLS genes (F8388_010207, F8388_010208) and two OAC genes (F8388_012844, F8388_012845) showed trichome-specific expression, while another OLS gene (F8388_010206) was enriched in trichomes and IMF (Table S7). Of the eight MEP pathway genes identified (Table S6), four were expressed in all samples, two showed trichome-specific expression—including HDS (F8388_013291) and HDR (F8388_013980)—and one was leaf-specific (Table S7). All six MEV pathway genes were classified as “expressed-in-all”. Notably, isopentenyl-diphosphate delta-isomerase (IDI), which interconverts the MEP- and MEV-derived IPP and DMAPP to form the isoprenoid backbone (Krause et al., 2023), was represented by two genes, one of which (F8388_016754) was trichome-specific. Together, these results indicate that genes from the polyketide and MEP pathways are preferentially expressed in trichomes. Importantly, no consistent expression differences were observed between hemp and marijuana varieties, nor between MF and FF, suggesting conservation of upstream cannabinoid biosynthetic pathways across chemotypes and sexual morphs.

**Figure 5.**
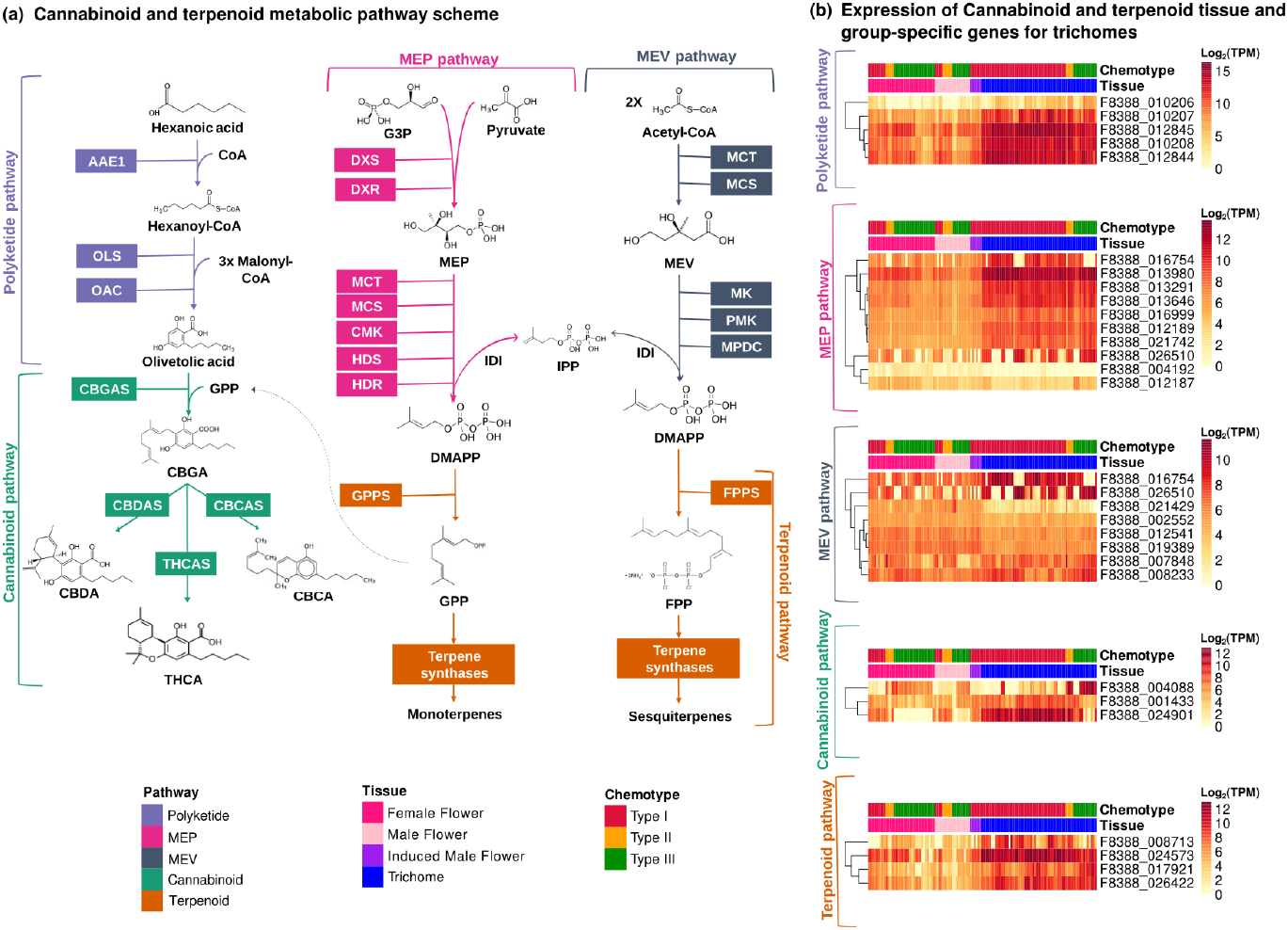
Cannabinoid and terpenoid biosynthesis. (a) Schematic representation of the metabolic pathways underlying cannabinoid and terpenoid production, with key enzymatic steps highlighted. (b) Expression profiles of cannabinoid- and terpenoid-related genes in trichome and whole-flower samples. Expression levels are shown across chemotypes to illustrate tissue specificity and chemotype-associated patterns.

We identified five genes directly involved in the cannabinoid biosynthetic pathway: three prenyltransferases (F8388_001433, F8388_008405, and F8388_013365), one Cannabidiolic Acid Synthase (CBDAS, F8388_004088), and one Tetrahydrocannabinolic Acid Synthase (THCAS, F8388_024901) (Table S6). Of these, THCAS and the prenyltransferase F8388_001433 were trichome-specific; CBDAS is group-enriched in FF and stems; the prenyltransferase F8388_008405 is mixed; and F8388_013365 is expressed-in-all tissues (Table S7). The classification of CBDAS as group-enriched in FF and stems, rather than trichome-specific, likely reflects both chemotype distribution and the methodology used by Barbosa-Xavier et al. (2024), which relies on median expression across samples. Approximately 70% of the trichome samples in the dataset originated from type I chemotypes, which typically show low CBDAS expression, potentially underestimating its trichome specificity. A similar bias could have occurred for THCAS had the dataset been skewed toward type III chemotypes. These observations highlight how sample composition and classification thresholds can affect the apparent tissue specificity of cannabinoid biosynthesis genes.

Cannabigerolic acid (CBGA), the central precursor of cannabinoids, is synthesized via alkylation of OAC and GPP, a reaction catalyzed by a CsPT1 aromatic prenyltransferase (CBGAS) (Innes & Vergara, 2023). The gene F8388_001433 shows high sequence identity to several prenyltransferases (Table S6), including CsPT7 (98.4%), CsPT1 (83.9%), and CsPT4 (78.7%). CsPT4 and CsPT7 are closely related to CsPT1, and CsPT4 has also been shown to catalyze CBGA formation in vivo (Luo et al., 2019; Rea et al., 2019). These similarities strongly suggest that F8388_001433 functions as an aromatic prenyltransferase involved in CBGA biosynthesis. Expression data support this functional assignment: THCAS, CBDAS, and CBGAS (F8388_001433) were more highly expressed in trichomes than in whole flowers (Figure 5b). Moreover, THCAS and CBGAS were preferentially expressed in type I and II chemotypes, while CBDAS was more highly expressed in type II and III chemotypes (Figure 5b), consistent with their roles in defining cannabinoid profiles.

Unexpected expression patterns were detected in certain type III hemp samples. Trichome samples SAMN09747685, SAMN09747688, and SAMN09747691, labeled as type III hemp (Finola strain) (Livingston et al., 2020), displayed type II-like expression profiles (Figure 5b). These samples are biological replicates from BioProject PRJNA483805, which otherwise shows consistent type III expression. We hypothesize that these samples may have been inadvertently contaminated with trichomes from type I varieties, such as Purple Kush or Hindu Kush, which were analyzed alongside Finola by Livingston et al. (2020) using two-photon microscopy and disc cell counting analysis. A second discrepancy was observed in the samples SAMN42101035, SAMN42101034, and SAMN42101033 from the MW6-15 cultivar (BioProject PRJNA1128734). Although described by Welling et al. (2023) as an industrial hemp line (type III), these samples displayed a type I expression profile, characterized by high THCAS expression and complete absence of CBDAS (Figure 5b). Such inconsistencies may reflect sample mislabeling, cross-contamination, or previously unrecognized chemotypic diversity, and warrant further validation. Regarding terpenoid biosynthesis, we identified eight candidate genes (Table S6): three with root-specific expression, three with trichome-specific, one group-enriched in trichomes and IMF, and one with low expression (Table S7). Trichome-specific genes (F8388_008713, F8388_024573, F8388_026422) and the group-enriched gene (F8388_017921) all exhibited markedly higher expression in trichomes compared to whole flower samples (Figure 5). Importantly, their expression showed no clear bias toward specific chemotypes. This pattern suggests that terpenoid biosynthesis is broadly active across both hemp and marijuana varieties, likely independent of cannabinoid chemotype-defining genes. Overall, despite chemotypic differences between hemp and marijuana, genes in the upstream cannabinoid and terpenoid pathways show similar expression profiles (Figure 5), highlighting conserved mechanisms of defense specialization in *C. sativa*.

## Conclusion

This study advances our understanding of *C. sativa* flower development by identifying OGs of MADS-box genes and analyzing their expression in MF and FF. Our results point to the involvement of specific MADS-box genes in flower organ development and sex differentiation. In parallel, sugar metabolism-related genes were linked to fertility and pollen wall formation, underscoring key metabolic processes in male reproductive structures. Expression profiling of cannabinoid and terpenoid biosynthetic genes further confirmed their strong tissue specificity, particularly in trichomes, emphasizing the finely tuned regulation of secondary metabolite production. Collectively, these findings establish a molecular framework for future functional studies and identify promising targets for metabolic engineering and breeding strategies aimed at optimizing flower traits and bioactive compound profiles in cannabis.

## Acknowledgements

This work was supported by Fundação Carlos Chagas Filho de Amparo à Pesquisa do Estado do Rio de Janeiro (FAPERJ), Coordenação de Aperfeiçoamento de Pessoal de Nível Superior-Brasil (CAPES; Finance Code 001), Conselho Nacional de Desenvolvimento Científico e Tecnológico (CNPq) and Programa de Apoio à Pesquisa, Inovação e Cultura (PAPIC - UENF). The funding agencies had no role in the design of the study and collection, analysis and interpretation of data and in writing.

## Funding Information

- Fundação Carlos Chagas Filho de Amparo à Pesquisa do Estado do Rio de Janeiro (FAPERJ);
- Coordenação de Aperfeiçoamento de Pessoal de Nível Superior-Brasil (CAPES; Finance Code 001);
- Conselho Nacional de Desenvolvimento Científico e Tecnológico (CNPq).
- Universidade Estadual do Norte Fluminense Darcy Ribeiro.

## Notes

### Competing Interest Statement

The authors have declared no competing interest.

