## Supplementary figures for "Transcription-based dissection of floral identity and trichome biosynthesis pathways in *Cannabis sativa* L"

Kevelin Barbosa-Xavier - https://orcid.org/0000-0002-3750-5331
Thiago Motta Venancio - https://orcid.org/0000-0002-2215-8082

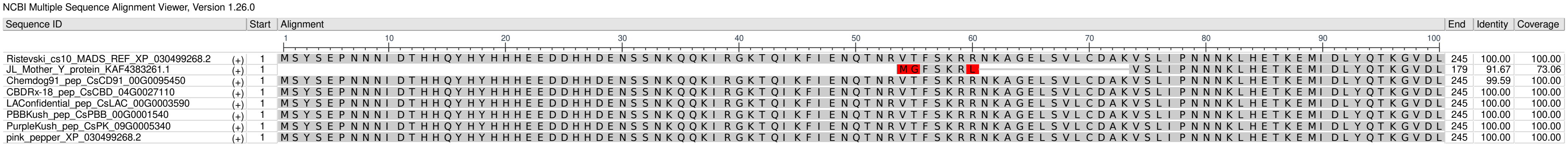


Figure S1 - Multiple Sequence Alignment Visualization of Clade AP3 from OG0000033

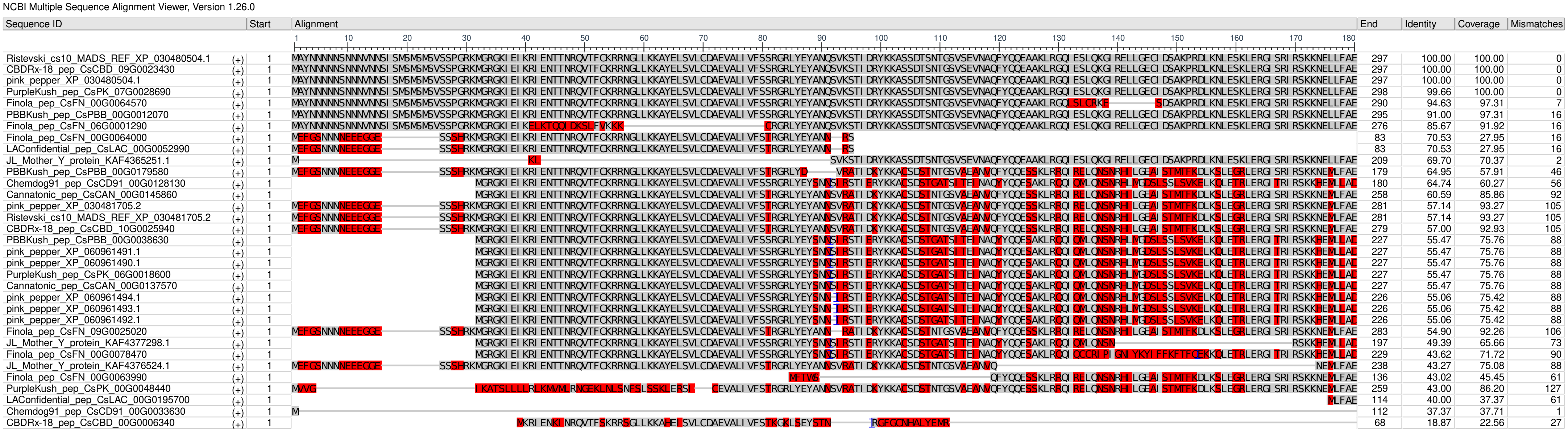


Figure S2 - Multiple Sequence Alignment Visualization of Clade AG from OG0000096

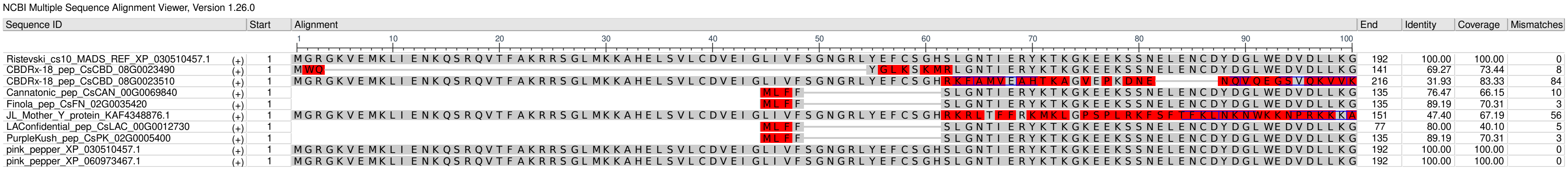


Figure S3 - Multiple Sequence Alignment Visualization of Clade FLC from OG0000096

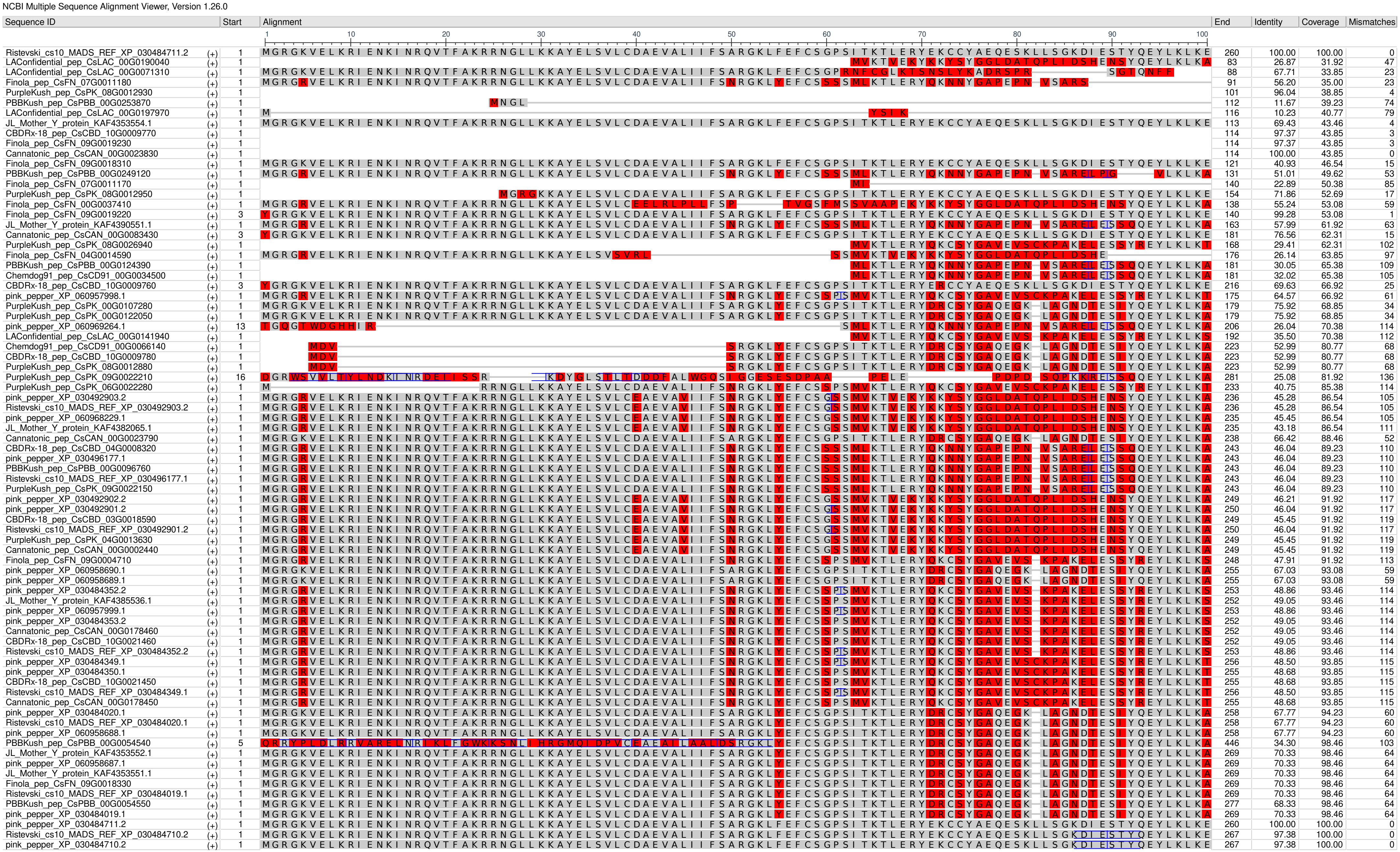


Figure S4 - Multiple Sequence Alignment Visualization of Clade SEP from OG0000096

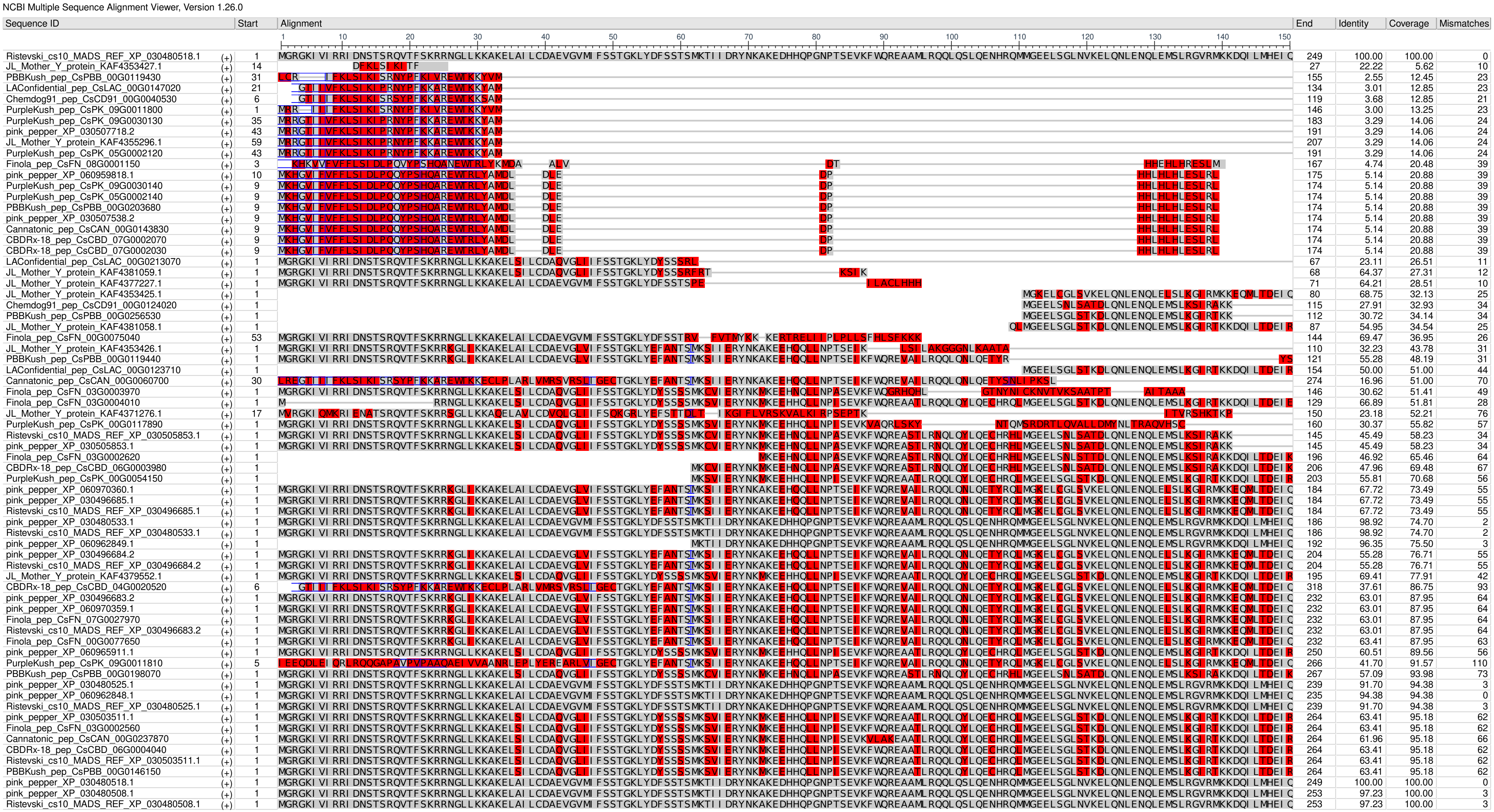


Figure S5 - Multiple Sequence Alignment Visualization of Clade AGL17 from OG0000207

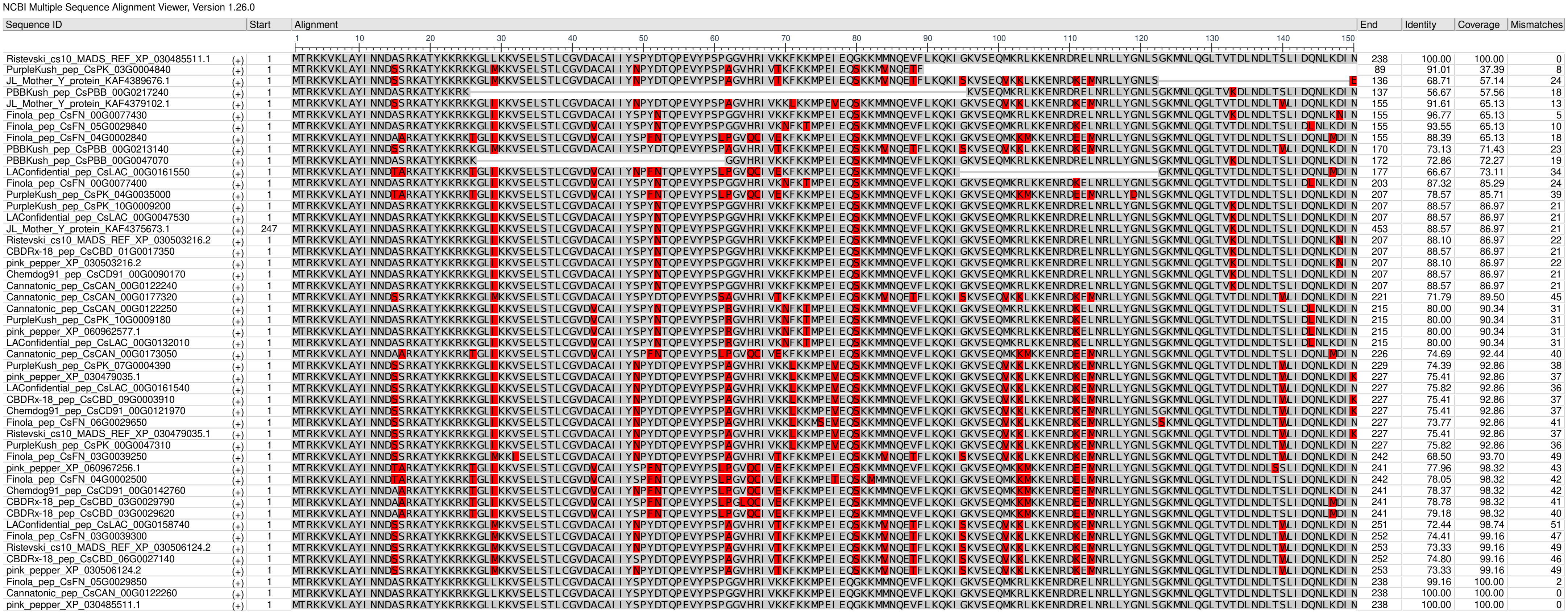


Figure S6 - Multiple Sequence Alignment Visualization of Clade Mg from OG0000366

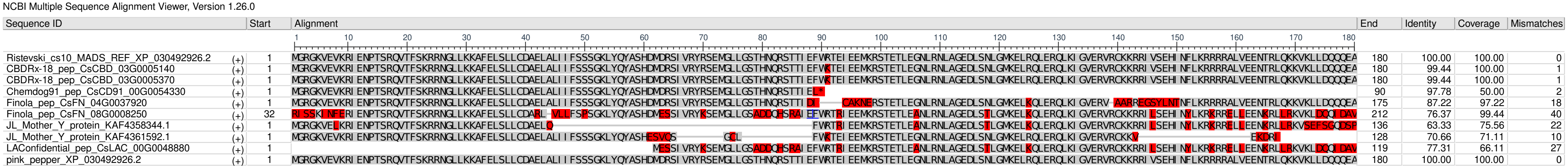


Figure S7 - Multiple Sequence Alignment Visualization of Clade FLC-like from OG0000433

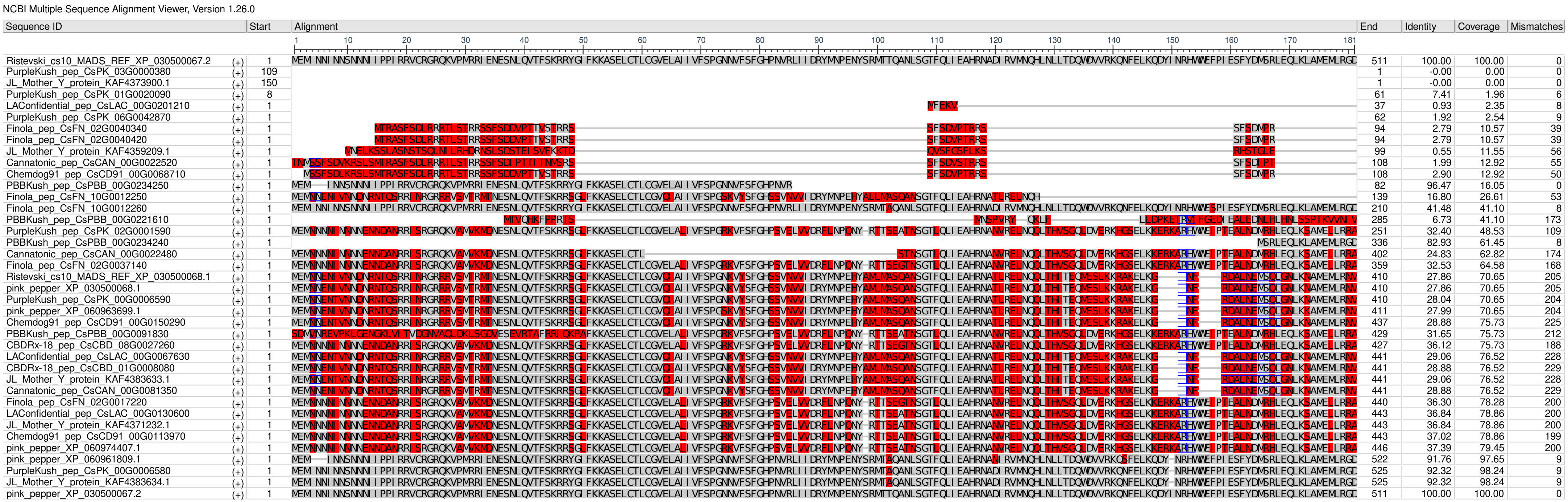


Figure S8 - Multiple Sequence Alignment Visualization of Clade Ma from OG0000580

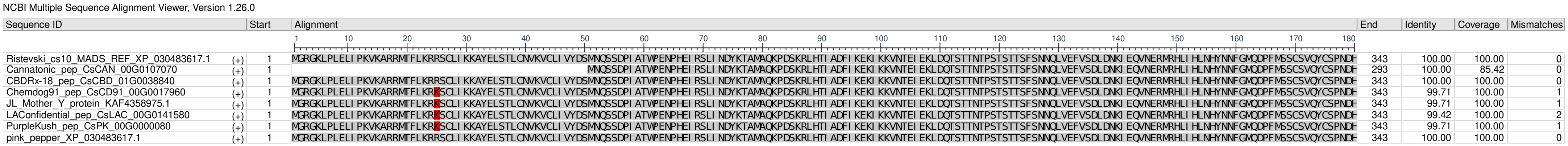


Figure S9 - Multiple Sequence Alignment Visualization of Clade Mb from OG0000736

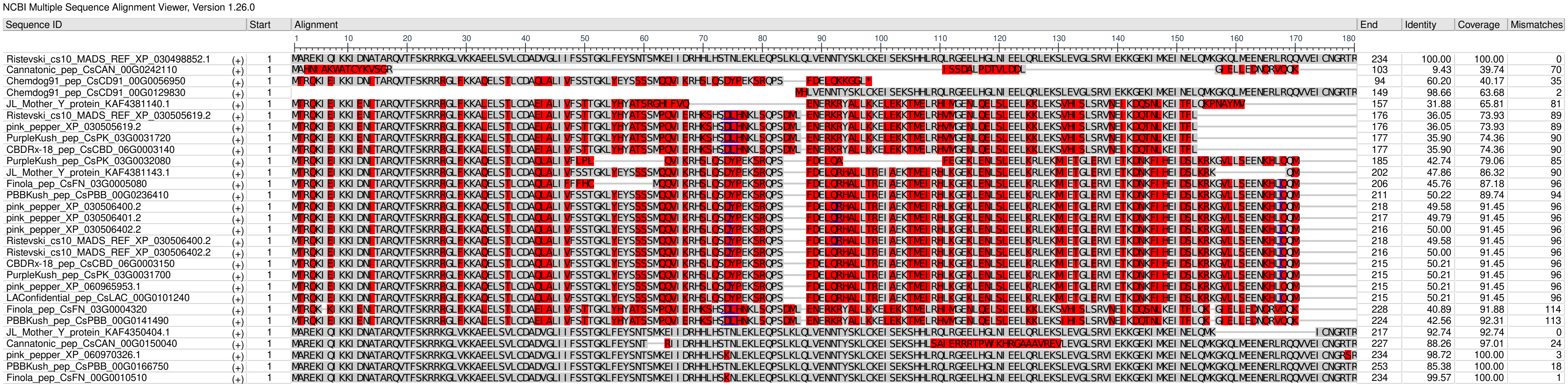


Figure S10 - Multiple Sequence Alignment Visualization of Clade SVP from OG0000955

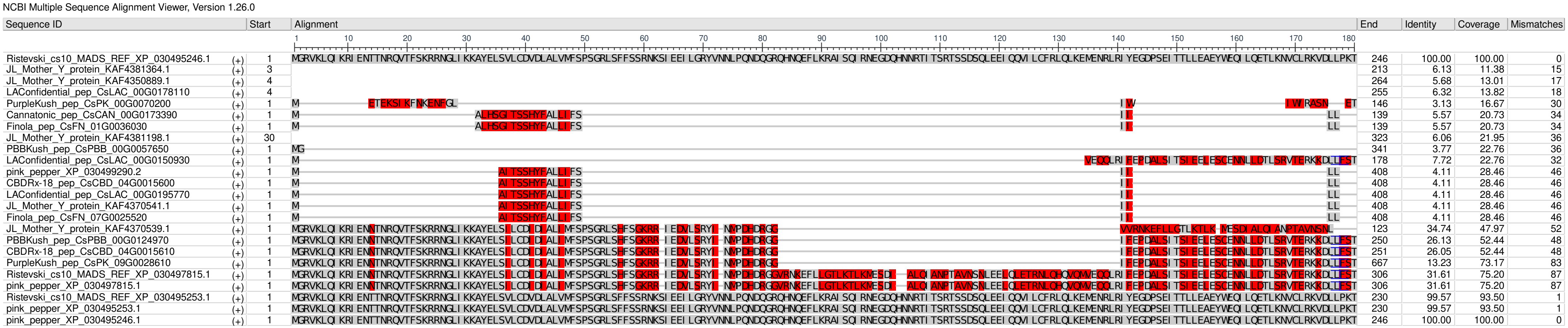


Figure S11 - Multiple Sequence Alignment Visualization of Clade MIKCS from OG0001105

NCBI Multiple Sequence Alignment Viewer, Version 1.26.0

| Sequence ID | | Start | Alignment | End | Identity | Coverage | Mismatches |
|---|---|---|---|---|---|---|---|
| Ristevski_cs10_MADS_REF_XP_030481490.1 | (+) | 1 | MGRGRVELKRIENKINRQVTFSKRRNGLLKKAHEISVLCDAEVAVIVFSHKGKLSEYSTDSCMEKILERYERYSAERQLVSMDSETQLQIMQGNWTVEYSRLTAKIELLQRNRRHYLGEDLDSLTLKDLTGLEQQIDISLKQIRARRNQLMNETISDLQKREKAAREENNLLEKKIKER | 254 | 100.00 | 100.00 | 0 |
| LAConfidential_pep_CsLAC_00G0219890 | (+) | 1 | M----------------------------------SFL---------------------------------------------------------------------------------------------------------------------------KAKAIQDRNNWTNQIKED | 103 | 9.23 | 33.86 | 61 |
| JL_Mother_Y_protein_KAF4351524.1 | (+) | 1 | MGRGRVEMKRIENKINRQVTFSKRRSGLLKKAHEISVLCDAEVALIVFSTKGKLSEYSTNSSMERILERYERYSAERQLMSNDPE-----QT---------------------------------------------------------------------DKALQEQNNILTKKIKEK | 188 | 40.45 | 68.90 | 67 |
| PBBKush_pep_CsPBB_00G0083660 | (+) | 1 | -------MKRIENKINRQVTFSKRRSGLLKKAHEISVLCDAEVALIVFSTKGKLSEYSTNSSMERILERYERYSAERQLMSNDPE-----QTGSWTLEHAKLKARMEVLQRNQKHFMGEELDSLSMKELQNLEQQLDSALKHVRSRKNQLMYESISDLQKKDKALQEQNNILTKKIKEK | 184 | 53.71 | 72.44 | 61 |
| PBBKush_pep_CsPBB_00G0054570 | (+) | 1 | MGRGRVELKRIENKINRQVTFSKRRNGLLKKAHEISVLCDAEVAVIVFSHKGKLSEYSTDSCMEKILERYERYSAERQLVSMDSETQLQIMQGNWTVEYSRLTAKIELLQRNRRHYLGEDLDSLTLKDLTGLEQQIDISLKQIRARRNQLMNETISDLQKREKAAREENNLLEKKMPV- | 185 | 75.11 | 72.83 | 7 |
| Chemdog91_pep_CsCD91_00G0095750 | (+) | 1 | --------------------------------------------------------------MEKILERYERYSAERQLVSMDSETQLQIMQGNWTVEYSRLTAKIELLQRNRRHYLGEDLDSLTLKDLTGLEQQIDISLKQIRARRNQLMNETISDLQKREKAAREENNLLEKKIKER | 192 | 100.00 | 75.59 | 0 |
| LAConfidential_pep_CsLAC_00G0100110 | (+) | 1 | --------------------------------------------------------------MEKILERYERYSAERQLVSMDSETQLQIMQGNWTVEYSRLTAKIELLQRNRRHYLGEDLDSLTLKDLTGLEQQIDISLKQIRARRNQLMNETISDLQKREKAAREENNLLEKKIKER | 192 | 100.00 | 75.59 | 0 |
| Cannatonic_pep_CsCAN_00G0023770 | (+) | 1 | MG------------------------------------------------RGKLSEYSTDSCMEKILERYERYSAERQLVSMDSETQLQIMQGNWTVEYSRLTAKIELLQRNRRHYLGEDLDSLTLKDLTGLEQQIDISLKQIRARRNQLMNETISDLQKREKAAREENNLLEKKIKER | 206 | 80.71 | 81.10 | 1 |
| JL_Mother_Y_protein_KAF4353550.1 | (+) | 1 | MGRGRVELKRIENKINRQVTFSKRRNGLLKKAHEISVLCDAEVAVIVFSHKGKLSEYSTDSCMEKILERYERYSAERQLVSMDSETQLQIMQGNWTVEYSRLTAKIELLQRNRRHYLGEDLDSLTLKDLTGLEQQIDISLKQIRARRNQLMNETISDLQKREKAAREENNLLEKKIALY | 218 | 82.68 | 85.83 | 8 |
| CBDRx-18_pep_CsCBD_00G0006380 | (+) | 1 | -------MKRIENKINRQVTFSKRRSGLLKKAHEISVLCDAEVALIVFSTKGKLSEYSTNSSMERILERYERYSAERQLMSNDPE-----QTGSWTLEHAKLKARMEVLQRNQKHFMGEELDSLSMKELQNLEQQLDSALKHVRSRKNQLMYESISDLQKKDKALQEQNNILTKKIKEK | 250 | 54.23 | 93.31 | 96 |
| Cannatonic_pep_CsCAN_00G0002570 | (+) | 1 | -------MKRIENKINRQVTFSKRRSGLLKKAHEISVLCDAEVALIVFSTKGKLSEYSTNSSMERILERYERYSAERQLMSNDPE-----QTGSWTLEHAKLKARMEVLQRNQKHFMGEELDSLSMKELQNLEQQLDSALKHVRSRKNQLMYESISDLQKKDKALQEQNNILTKKIKEK | 250 | 54.23 | 93.31 | 96 |
| Finola_pep_CsFN_09G0004890 | (+) | 1 | MGRGRVEMKRIENKISRQVTFSKRRSGLLKKAHEISVLCDAHVALIVFSTKGKLFNFSSSSSMEEILERYESYSTNAQKHLQAAPTTSESQDSSKWTFEFPKLMSRVEILQKNLRNCMGEDLALLSMRELQNIEQQIDIGLKRVRTRKNQIMHETLSDLQKKAKAIQDRNNWTNQIKED | 261 | 43.38 | 95.67 | 125 |
| pink_pepper_XP_030485101.2 | (+) | 1 | MGRGRVEMKRIENKISRQVTFSKRRSGLLKKAHEISVLCDAHVALIVFSTKGKLFNFSSSSSMEEILERYESYSTNAQKHLQAAPTTSESQDSSKWTFEFPKLMSRVEILQKNLRNCMGEDLALLSMRELQNIEQQIDVGLKRVRTRKNQIMHETLSDLQKKAKAIQDRNNWTNQIKED | 262 | 42.86 | 95.67 | 126 |
| Ristevski_cs10_MADS_REF_XP_030485101.2 | (+) | 1 | MGRGRVEMKRIENKISRQVTFSKRRSGLLKKAHEISVLCDAHVALIVFSTKGKLFNFSSSSSMEEILERYESYSTNAQKHLQAAPTTSESQDSSKWTFEFPKLMSRVEILQKNLRNCMGEDLALLSMRELQNIEQQIDVGLKRVRTRKNQIMHETLSDLQKKAKAIQDRNNWTNQIKED | 262 | 42.86 | 95.67 | 126 |
| PurpleKush_pep_CsPK_08G0026950 | (+) | 1 | MGRGRVEMKRIENKISRQVTFSKRRSGLLKKAHEISVLCDAHVALIVFSTKGKLFNFSSSSSMEEILERYESYSTNAQKHLQAAPTTSESQDSSKWTFEFPKLMSRVEILQKNLRNCMGEDLALLSMRELQNIEQQIDVGLKRVRTRKNQIMHETLSDLQKKAKAIQDRNNWTNQIKED | 261 | 43.01 | 95.67 | 126 |
| PurpleKush_pep_CsPK_06G0022270 | (+) | 1 | MGRGRVEMKRIENKISRQVTFSKRRSGLLKKAHEISVLCDAHVALIVFSTKGKLFNFSSSSSMEEILERYESYSTNEQKHLQAAPTTSESQDSSKWTFEFPKLMSRVEILQKNLRNCMGEDLALLSMRELQNIEQQIDIGLKRVRTRKNQIMHETLSDLQKKAKAIQDRNNWTNQIKED | 261 | 43.75 | 95.67 | 124 |
| PBBKush_pep_CsPBB_00G0195530 | (+) | 1 | MGRGRVEMKRIENKISRQVTFSKRRSGLLKKAHEISVLCDAHVALIVFSTKGKLFNFSSSSSMEEILERYESYSTNEQKHLQAAPTTSESQDSSKWTFEFPKLMSRVEILQKNLRNCMGEDLALLSMRELQNIEQQIDIGLKRVRTRKNQIMHETLSDLQKKAKAIQDRNNWTNQIKED | 261 | 43.75 | 95.67 | 124 |
| CBDRx-18_pep_CsCBD_10G0021440 | (+) | 1 | MGRGRVEMKRIENKISRQVTFSKRRSGLLKKAHEISVLCDAHVALIVFSTKGKLFNFSSSSSMEEILERYESYSTNAQKHLQAAPTTSESQDSSKWTFEFPKLMSRVEILQKNLRNCMGEDLALLSMRELQNIEQQIDIGLKRVRTRKNQIMHETLSDLQKKAKAIQDRNNWTNQIKED | 261 | 43.38 | 95.67 | 125 |
| JL_Mother_Y_protein_KAF4385535.1 | (+) | 1 | MGRGRVEMKRIENKISRQVTFSKRRSGLLKKAHEISVLCDAHVALIVFSTKGKLFNFSSSSSMEEILERYESYSTNEQKHLQAAPTTSESQDSSKWTFEFPKLMSRVEILQKNLRNCMGEDLALLSMRELQNIEQQIDIGLKRVRTRKNQIMHETLSDLQKKAKAIQDRNNWTNQIKED | 261 | 43.75 | 95.67 | 124 |
| Finola_pep_CsFN_04G0023150 | (+) | 1 | MGRGRVEMKRIENKINRQVTFSKRRSGLLKKAHEISVLCSSSPSRSTLPIPGNVSIFDVDDSMERILERYERYSAERQLMSNDPE-----QTGSWTLEHAKLKARMEVLQRNQKHFMGEELDSLSMKELQNLEQQLDSALKHVRSRKNQLMYESISDLQKKDKALQEQNNILTKKIKEK | 273 | 46.64 | 96.06 | 112 |
| Finola_pep_CsFN_04G0023620 | (+) | 1 | MYRHRTAEKKYQDYLYRKVTFSKRRSGLLKKAHEISVLCDAEVALLVFSTKGKLSEYSTNSSMERILERYERYSAERQLMSNDPE-----QTGSWTLEHAKLKARMEVLQRNQKHFMGEELDSLSMKELQNLEQQLDSALKHVRSRKNQLMYESISDLQKKDKALQEQNNILTKKIKEK | 391 | 33.67 | 96.06 | 109 |
| Ristevski_cs10_MADS_REF_XP_030485608.2 | (+) | 1 | MGRGRVEMKRIENKINRQVTFSKRRSGLLKKAHEISVLCDAEVALLVFSTKGKLSEYSTNSSMERILERYERYSAERQLMSNDPE-----QTGSWTLEHAKLKARMEVLQRNQKHFMGEELDSLSMKELQNLEQQLDSALKHVRSRKNQLMYESISDLQKKDKALQEQNNILTKKIKEK | 256 | 55.26 | 96.06 | 97 |
| pink_pepper_XP_030485608.2 | (+) | 1 | MGRGRVEMKRIENKINRQVTFSKRRSGLLKKAHEISVLCDAEVALLVFSTKGKLSEYSTNSSMERILERYERYSAERQLMSNDPE-----QTGSWTLEHAKLKARMEVLQRNQKHFMGEELDSLSMKELQNLEQQLDSALKHVRSRKNQLMYESISDLQKKDKALQEQNNILTKKIKEK | 256 | 55.26 | 96.06 | 97 |
| pink_pepper_XP_060968230.1 | (+) | 1 | MGRGRVEMKRIENKINRQVTFSKRRSGLLKKAHEISVLCDAEVALLVFSTKGKLSEYSTNSSMERILERYERYSAERQLMSNDPE-----QTGSWTLEHAKLKARMEVLQRNQKHFMGEELDSLSMKELQNLEQQLDSALKHVRSRKNQLMYESISDLQKKDKALQEQNNILTKKIKEK | 256 | 55.26 | 96.06 | 97 |
| pink_pepper_XP_030481491.1 | (+) | 1 | MGRGRVELKRIENKINRQVTFSKRRNGLLKKAHEISVLCDAEVAVIVFSHKGKLSEYSTDSCMEKILERYERYSAERQLVSMDSETQL---QGNWTVEYSRLTAKIELLQRNRRHYLGEDLDSLTLKDLTGLEQQIDISLKQIRARRNQLMNETISDLQKREKAAREENNLLEKKIKER | 251 | 98.82 | 98.82 | 0 |
| PurpleKush_pep_CsPK_08G0012860 | (+) | 1 | MGRGRVELKRIENKINRQVTFSKRRNGLLKKAHEISVLCDAEVAVIVFSHKGKLSEYSTDSCMEKILERYERYSAERQLVSMDSETQLQIMQGNWTVEYSRLTAKIELLQRNRRHYLGEDLDSLTLKDLTGLEQQIDISLKQIRARRNQLMNETISDLQKREKAAREENNLLEKKIKER | 254 | 100.00 | 100.00 | 0 |
| Finola_pep_CsFN_09G0018350 | (+) | 1 | MGRGRVELKRIENKINRQVTFSKRRNGLLKKAHEISVLCDAEVAVIVFSHKGKLSEYSTDSCMEKILERYERYSAERQLVSMDSETQLQIMQGNWTVEYSRLTAKIELLQRNRRHYLGEDLDSLTLKDLTGLEQQIDISLKQIRARRNQLMNETISDLQKREKAAREENNLLEKKIKER | 254 | 99.61 | 100.00 | 1 |

Figure S12 - Multiple Sequence Alignment Visualization of Clade SQUA from OG0001251

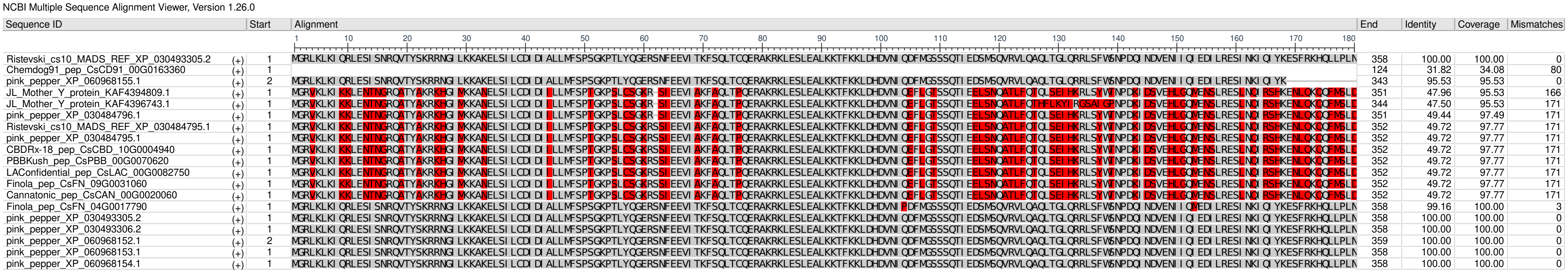


Figure S13 - Multiple Sequence Alignment Visualization of Clade MIKCS from OG0001495

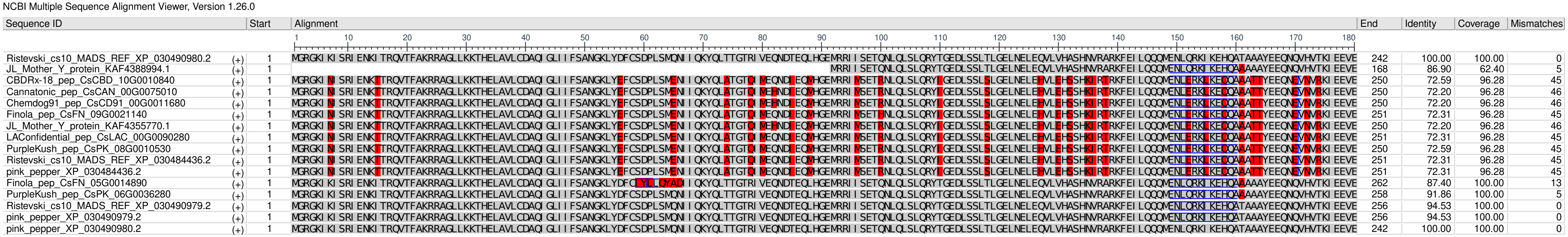


Figure S14 - Multiple Sequence Alignment Visualization of Clade Bsister from OG0002194

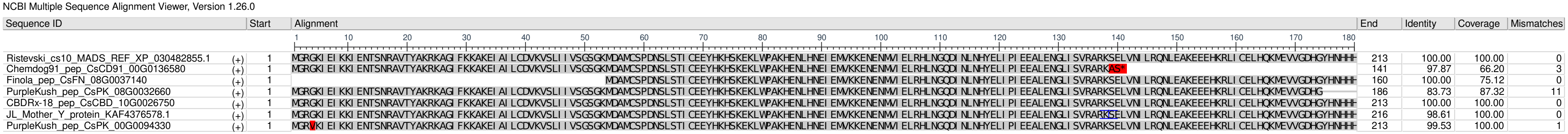


Figure S15 - Multiple Sequence Alignment Visualization of Clade PI/GLO from OG0005875

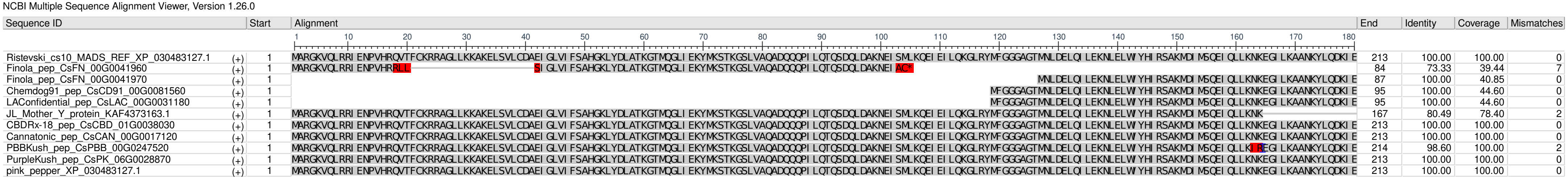


Figure S16 - Multiple Sequence Alignment Visualization of Clade AGL12 from OG0006023

NCBI Multiple Sequence Alignment Viewer, Version 1.26.0

| Sequence ID | | Start | Alignment | End | Identity | Coverage | Mismatches |
|---|---|---|---|---|---|---|---|
| Ristevski_cs10_MADS_REF_XP_030494537.1 | (+) | 1 | MGRGKIEIKRIENANSRQVTFSKRRAGLLKKAQELAILCDAEVAVIIFSNTGKLFEYSSSGMKRTLARYNKCVDYPEGAIVDKTEKQDSKEVDDLKDEIAKLQRKQSRLLGEDLSGMSSKELQHLEHQLNDGLISVKERKEKLLKEQLEQSRLQEQRAVLENETLRRQIEELRCLFPRAD | 254 | 100.00 | 100.00 | 0 |
| LAConfidential_pep_CsLAC_00G0085810 | (+) | 1 | MLY IVSLILSNF SVFQEQRAVLENETLRRQIEELRCLFPRAD | 117 | 41.57 | 45.67 | 10 |
| PurpleKush_pep_CsPK_00G0018340 | (+) | 1 | MGRGKIEIKRIENANSRQVTFSKRRAGLLKKAQELAILCDAEVAVIIFSNTGKLFEYSSSGMKRTLARYNKCVDYPEGAIVDKTEKQDSKEVDDLKDEIAKLQRKQSRLLGEDLSGMSSKELQHLEHQLNDGLISVKERKEKLLKEQLEQSRLQEQRAVLENETLRRQIEELRCLFPRAD | 225 | 77.47 | 88.19 | 28 |
| Chemdog91_pep_CsCD91_00G0161240 | (+) | 1 | MGRGKIEIKRIENANSRQVTFSKRRAGLLKKAQELAILCDAEVAVIIFSNTGKLFEYSSSGMKRTLARYNKCVDYPEGAIVDKTEKQDSKEVDDLKDEIAKLQRKQSRLLGEDLSGMSSKELQHLEHQLNDGLISVKERKEKLLKEQLEQSRLQEQRAVLENETLRRQIEELRCLFPRAD | 254 | 100.00 | 100.00 | 0 |
| PurpleKush_pep_CsPK_04G0000610 | (+) | 1 | MQKGKIEIKRIENANSRQVTFSKRRAGLLKKAQELAILCDAEVAVIIFSNTGKLFEYSSSGMKRTLARYNKCVDYPEGAIVDKTEKQDSKEVDDLKDEIAKLQRKQSRLLGEDLSGMSSKELQHLEHQLNDGLISVKERKEKLLKEQLEQSRLQEQRAVLENETLRRQIEELRCLFPRAD | 264 | 95.45 | 100.00 | 2 |

Figure S17- Multiple Sequence Alignment Visualization of Clade AGL15 from OG0007459
